# Microbiome determinants of productivity in whiteleg shrimp aquaculture

**DOI:** 10.1101/2024.09.20.614022

**Authors:** Xiaoyu Shan, Kunying Li, Patrizia Stadler, Martha Borbor, Guillermo Reyes, Ramiro Solórzano, Esmeralda Chamorro, Bonny Bayot, Otto X. Cordero

## Abstract

Aquaculture holds immense promise for addressing the food needs of our growing global population. Yet, a quantitative understanding of the factors that control its efficiency and productivity has remained elusive. In this study, we address this knowledge gap by focusing on the microbiome determinants of productivity, more specifically animal survival and growth, for one of the most predominant animal species in global aquaculture, whiteleg shrimp (*Penaeus vannamei*). Through analysis of the shrimp-associated microbiome from 610 aquaculture samples across Asia and Latin America, we established the presence of core phylogenetic groups, widely prevalent across aquaculture conditions in disparate geographic locations and including both pathogenic and beneficial microbes. Focusing on early stages of growth (larval hatcheries), we showed that the composition of the microbiome alone can predict approximately 50% of the variation in shrimp larvae survival rates. Taxa associated with high survival rates share recently acquired genes that appear to be specific to aquaculture conditions. These genes are involved in the biosynthesis of growth factors and in protein degradation, underscoring the potential role of beneficial microorganisms in nutrient assimilation. In contrast, the predictability of the microbiome on the adult shrimp weight in grow-out farms is weaker (10-20%), akin to observations in the context of livestock. In conclusion, our study unveils a novel avenue for predicting productivity in shrimp aquaculture based on microbiome analysis. This paves the way for targeted manipulation of the microbiome as a strategic approach to enhancing aquaculture efficiency from the earliest developmental stages.

## Introduction

In aquaculture, and animal farming in general, animal survival and individual growth yield are key factors that affect global measures of performance^1–3^. Improving resilience against disease (i.e. survival) and the efficiency with which animals assimilate their feed (individual growth yield) lies at the heart of addressing the twin challenges faced by the future food systems: elevating production to satisfy the escalating global food demand while reducing environmental impacts to align with sustainable development goals. Within this framework, the host microbiome is widely acknowledged to play an important role^4–9^. Beneficial microbes have the potential to improve survival and growth by facilitating the digestion and assimilation of nutrients^10^ or safeguarding against diseases^11^. Conversely, pathogenic microbes may compromise production by hindering growth or increasing mortality^12^, thus reducing yield in relation to resource consumption.

Since the late 1980s, aquaculture witnessed a rapid increase, in contrast to the stable production of marine capture fisheries, providing high-quality edible proteins and setting a new record with earnings of USD 265 billion (only for aquatic animals) and employing 20.7 million people as of 2020^13^. Whiteleg shrimp (*Penaeus vannamei*) farming, in particular, leads in aquaculture animal production with an annual production of 5.8 million tonnes in 2020^13^. Despite its rapid expansion, productivity of shrimp aquaculture frequently faces setbacks due to feed wastage^14^ and pathogen infections^15^, underscoring the need for technological advancements to improve production through microbiome engineering. However, the variable outcomes associated with probiotic use in shrimp farming have generated both enthusiasm and skepticism^16,17^. This highlights a significant gap in our understanding caused by a lack of comprehensive, systematic studies on how the microbiome composition affects productivity in shrimp aquaculture. Addressing this gap is crucial for leveraging microbiome-based strategies to their fullest potential.

In this study, we investigate the linkage between the microbiome and production variables in the aquaculture of *P. vannamei*. We start by characterizing a shrimp-associated microbiome across hatcheries and farms distributed across the globe. Subsequently, we focus separately on the two major phases of shrimp production: the hatchery phase and the grow-out phase. In the hatchery phase where shrimp transition from nauplii to postlarvae, survival rates are critical for productivity as early-stage shrimp are particularly vulnerable to environmental stressors and pathogenic threats. We investigate how the microbiome influences these survival rates, taking advantage of the tightly controlled abiotic conditions in hatcheries. In the grow-out phase, where the major goal is achieving optimal growth as shrimp advance to marketable sizes, we focus on body weight of shrimp as a direct indicator of growth efficiency and productivity under varying conditions of water quality and nutrient availability. Through this approach, we aim to shed light on the potential of microbiome interventions in enhancing the sustainability and productivity of shrimp aquaculture.

## Results

### Characterizing the global shrimp-associated microbiome

To better understand the role of the microbiome on shrimp production, we first compiled a global dataset of aquaculture shrimp-associated microbiome (Method). Briefly, we re-analyzed microbiome sequencing data of 610 samples^18–27^ from 12 geographically distant shrimp cultures in Asia and Latin America, including 232 samples of shrimp larvae microbiome and 378 samples of intestine (gut) microbiome from juvenile or adult shrimp (Figure 1A, Table S1). Recognizing the coastal ocean as the natural source of microbes in marine aquaculture, we also incorporated 154 samples of coastal seawater microbiome^28–30^. This extensive effort yielded a database of 56,513 ASVs (Amplicon Sequence Variants), including 48,514 shrimp-associated ASVs and 7,999 coastal seawater ASVs, now publicly available for further aquaculture research.

**Figure 1.**
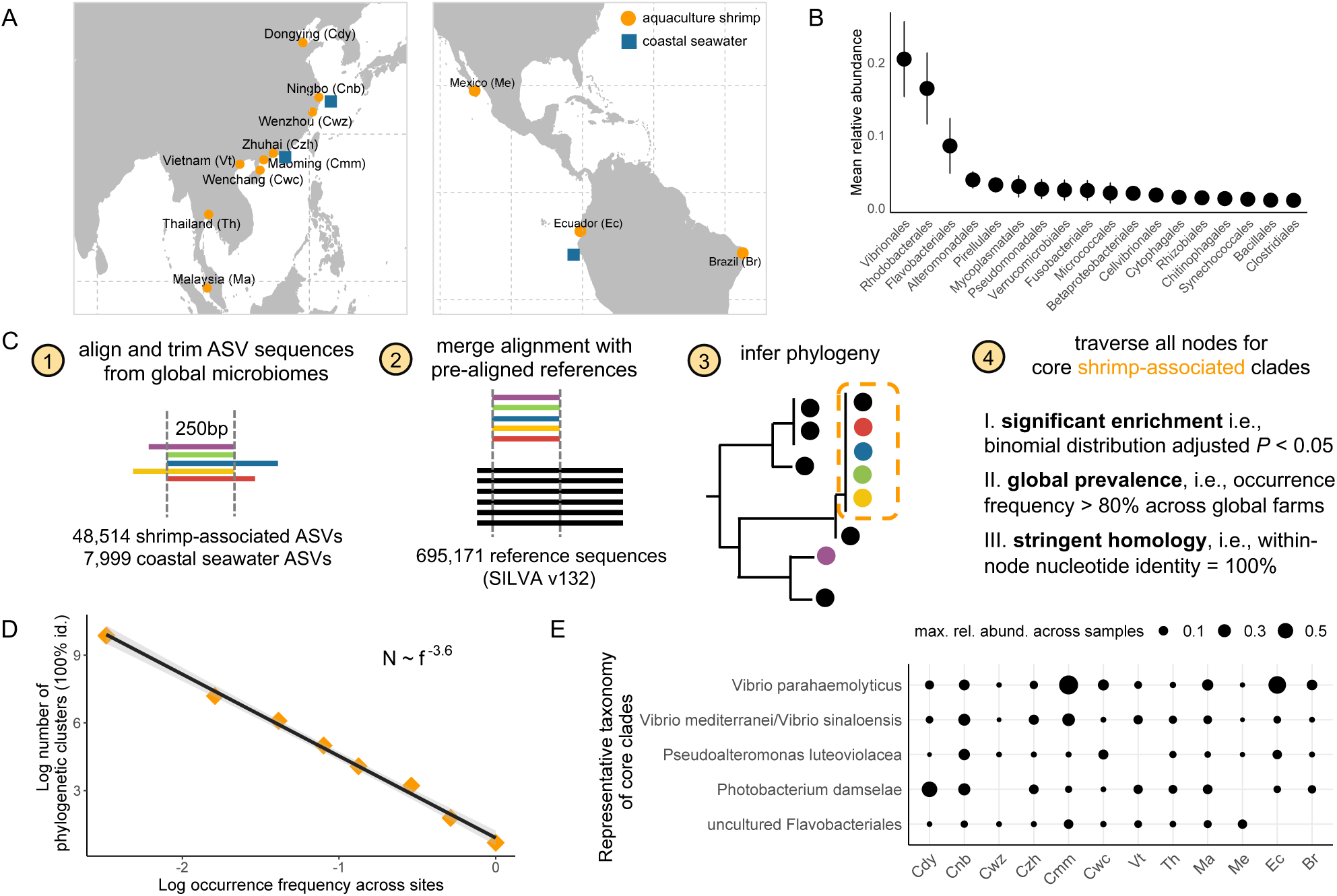
Identifying core clades of shrimp-associated microbiome from global aquaculture. (A) Geographic distribution of microbiome datasets compiled in this study. Orange circle indicates shrimp-associated microbiome. Blue square indicates coastal seawater microbiome. (B) Ranked mean relative abundance of bacterial taxa across global shrimp-associated microbiomes. (C) A bioinformatic pipeline is developed to identify core clades of shrimp-associated microbiome. See Methods for full descriptions. (D) A power law relationship between the number of phylogenetic clusters N and the occurrence frequency of phylogenetic clusters f. (E) Representative taxonomy of the identified core clades of global shrimp-associated microbiome. Size of dots indicates maximal relative abundance of a clade in a microbiome.

With this dataset of global shrimp-associated microbiome, we begin by asking whether the aquaculture environment selects for specific taxonomic clades, or it simply reflects a random assortment from the surrounding seawater. We find that the mean pairwise phylogenetic distances (MPD) between shrimp-associated sequences and coastal seawater sequences are significantly higher than that expected by a null model (Nearest Relatedness Index = 9.7, *P* < 0.001, see Method), suggesting a selective adaptation of specific microbes to a farm-associated environment. Indeed, we identified taxonomic groups that are significantly enriched or diluted in shrimp-associated microbiome. Oligotrophic taxa such as SAR11 and SAR86 are significantly less prevalent (Figure S1, adjusted *P* < 0.01), while copiotrophic taxa such as Vibrionales, Rhodobacterales and Flavobacteriales are among the most abundant ones across global aquaculture shrimp microbiome (Figure 1B). Interestingly, these taxa were also among the most abundant taxa in microbial communities assembled on marine particles of polysaccharides such as chitin^31,32^. Vibrionales are known for fast growth rate, especially in saline and estuarine environments (0.5∼3% NaCl) and high temperature (25∼30°C) featured by aquaculture conditions^33^. Flavobacteriales are noted for their capabilities in degrading high-molecular-weight polysaccharide and proteins^34,35^, whereas Rhodobacterales are specialized in scavenging metabolic byproducts such as organic acids and amino acids^34,35^.

Having elucidated the overall taxonomic composition of shrimp-associated microbiome, we further ask if there are core clades at finer phylogenetic scale that are conservative and significantly enriched across global shrimp cultures (Figure 1C, see Method). By grouping ASV ribotypes into phylogenetic clusters, we identify a power law relationship between the number of phylogenetic clusters (*N*) and their occurrence frequency (*f*): 𝑁∼𝑓^−*a*^, with 𝑎 = −3.6 (Figure 1D). This power law exponent is larger in magnitude than those observed in other systems^36^, including the global ocean microbiome (where 𝑎 = -1.7)^37^. This indicates a more rapid decline in the occurrence frequency as we move from the most to the least ubiquitous cluster, suggesting a highly uneven distribution of phylogenetic clusters across different shrimp farms. Nonetheless, we were able to identify five phylogenetic clades that are significantly enriched across shrimp cultures worldwide (Figure 1E). This includes three clades related to Vibrionales, represented by *Vibrio parahaemolyticus*, *Vibrio mediterranei*/*sinaloensis* and *Photobacterium damselae*, all of which are commonly known as aquaculture pathogens of fish or shrimp^38–40^. In particular, *V. parahaemolyticus* (Vp), the causative agent of acute hepatopancreatic necrosis disease (AHPND), is notorious for causing enormous economic losses to global shrimp farms every year^12^. In addition to these pathogens, a clade in the order of Alteromonadales, represented by *Pseudoalteromonas luteoviolaceae*, is also part of the core shrimp farm microbiome. Members of this clade have been shown to inhibit shrimp pathogens by producing antimicrobial compounds^41^, suggesting that some of the core microbiome members may confer beneficial effects to the host.

### Microbiome composition predicts survival in shrimp larvae hatcheries

We first probe into the impacts of microbiome composition on productivity at the hatchery stage. Here, shrimp larvae are nurtured from their earliest nauplii (N) stage to the postlarvae (PL) phase, preparing them for transfer to large-scale farms. We tracked the microbiome succession through developmental stages (nauplii, zoea, mysis and postlarvae) in a shrimp hatchery over a period of 18 days (Figure 2A, see Method). Initially, the microbiome composition of the larvae is most similar to the surrounding seawater, especially during the early nauplii and zoea stages (Figure 2B). This resemblance gradually diminishes as the larvae mature towards the PL stage (from ∼30% shared ASVs to ∼8% shared ASVs, ∼ 20% compositional similarity to ∼ 10% compositional similarity). At this stage, the larvae-associated microbiome became most similar to that of the larval feed, which was based on the aquatic crustacean *Artemia franciscana* (∼ 40% shared ASVs, ∼ 30% compositional similarity; Figure 2B, Figure S2). Such opposite trends of microbiomes similarities with seawater and feed indicates a transition of microbial influences from environmental to dietary sources. Interestingly, the bacteria administered through probiotics (> 80% composed of Lactobacillaceae) appear to be unable to colonize the host, never reaching relative abundance above 1% throughout all larval developmental stages. This result highlights the need to develop new microbiome therapies based on microorganisms native to the conditions of seawater aquaculture.

**Figure 2.**
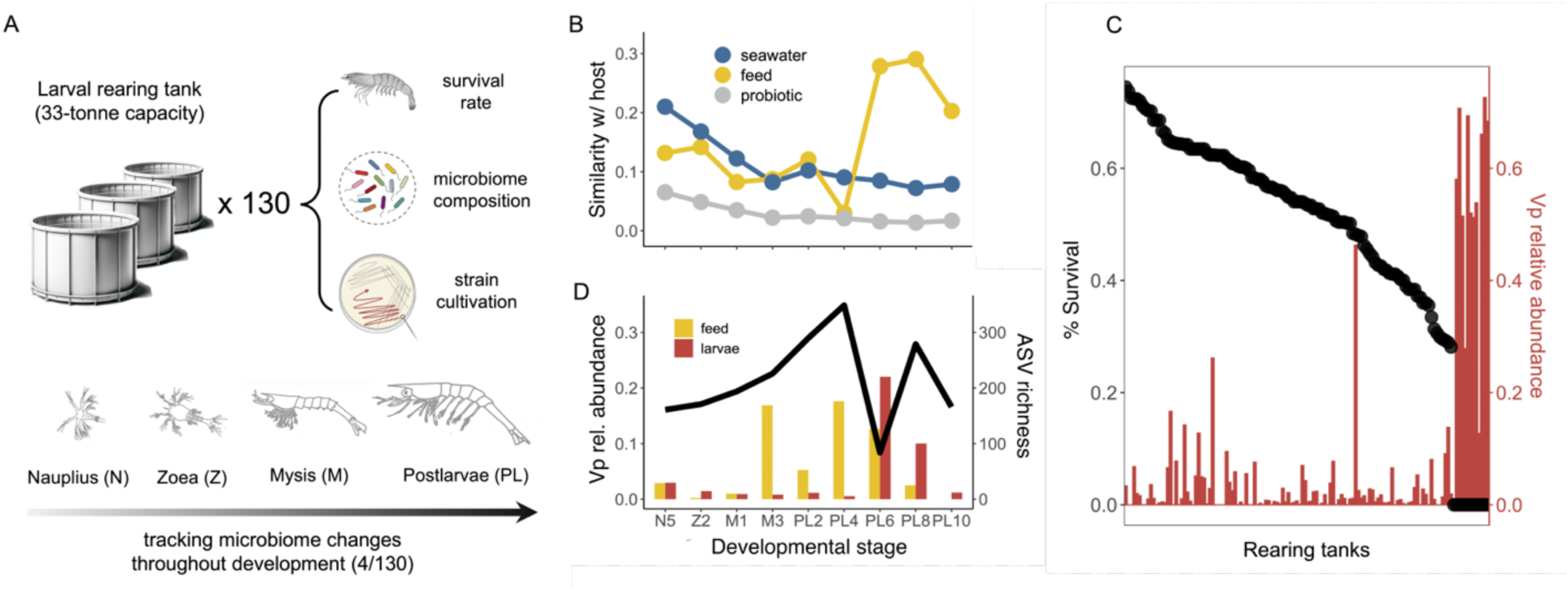
Microbiome characterization in a shrimp hatchery. (A) Shrimp larvae were sampled from 130 33-tonne capacity tanks where the survival rate of shrimp larvae was counted and recorded. The sampled shrimp larvae were further used for microbiome sequencing and bacterial strain isolation. For four selected tanks, we sampled throughout the developmental stages from nauplius, zoea, mysis to postlarvae. The potential sources of microbiome such as seawater, feed (microalgae at the nauplius and zoea stage and *Artemia* at the mysis and postlarvae stage) as well as probiotics were also sampled and sequenced. (B) Similarity between shrimp larvae-associated microbiome composition and microbiome composition of potential sources. Similarity is calculated as 1-Bray-Curtis dissimilarity based on relative abundance. (C) Black dots show the ranked survival rate of shrimp larvae across 130 tanks. Red bar shows relative abundance of *V. parahaemolyticus* in the corresponding shrimp larvae-associated microbiome. (D) Black line indicates richness of shrimp larvae-associated microbiome across developmental stages. Red bar and yellow bar indicate relative abundances of *V. parahaemolyticus* across developmental stages in shrimp larvae microbiome and microbiome in the feed.

To map microbiome composition to productivity, we sampled extensively across 130 tanks of 33-tonne capacity from this hatchery (Figure 2A, see Method). These tanks represent well-controlled replicate environments that undergo identical feeding regimes, therefore the differences in animal survival represents a major cause of variability in productivity and feed efficiency. Consistent with observations of Vp causing large scale losses in production, we found in this hatchery that 10% (13 out of 130) of tanks showed signs of collapse (no harvest) and that the microbiome of the larvae in these tanks was made of over 50% Vp (Figure 2C). Our microbiome development data showed that the rise of Vp in the larval microbiome was a sudden event, with an increase from nearly zero to ∼ 20% relative abundance from day 4 to day 6 at the PL stage (Figure 2D). The observed surge in Vp abundance first appeared in the feed (i.e., Artemia) before being mirrored in the larvae (Figure 2D, Figure S3), aligning with the developmental phase when the larvae microbiome is primarily shaped by their feed rather than the ambient seawater. This pattern indicates that *Artemia* can be a primary vector for Vp transmission. Interestingly, excluding these collapsed tanks, the remaining 90% of tanks had variable larvae survival rates, from ∼30% to ∼80% (Figure 2C). These rates were only poorly correlated to Vp’s abundance (Pearson’s r = 0.007, *P* = 0.98), indicating that other microbiome components may instead determine productivity.

Given this large variation in larvae survival across tanks, we asked to what extent it can be attributed to differences in microbiome composition. To address this question, we employed a Random Forest model to predict the survival rate of shrimp larvae based on their microbiome composition. We utilized a leave-one-out (LOO) cross-validation approach, allowing us to maximize the use of all available samples while ensuring an independent test set to prevent overfitting. With all samples, we find that microbiome composition can predict nearly 70% of the observed variation in larvae survival (LOO-R^2^ = 0.68, Figure 3A), which is as expected since a single ASV of Vp is known to be a strong indictor. We then ask how predictable the survival rate is if we exclude all the collapsed tanks. Remarkably, we found that microbiome composition alone still predicts nearly 50% of variation in survival rate (LOO-R^2^ = 0.46, Figure 3B). To test the robustness of this prediction, we conducted a series of random tests by reshuffling the survival rates across tanks 100 times and reassessing predictability of the model. These randomization tests consistently resulted in a significantly reduced predictive power (average LOO-R^2^ < 0.05, Figure 3A), demonstrating that specific microbiome compositions are indeed closely linked to survival outcomes in shrimp larvae, instead of due to chance. We further show that the prediction power of the model linearly increases with the number of samples (2.5% net increase in LOO-R^2^ per 10 samples added, linear regression R^2^ = 0.94, *P* < 0.001, Figure 3C), suggesting that the predictability of larvae survival by microbiome composition can be further elevated by expanding the sample space (i.e., sampling more tanks).

**Figure 3.**
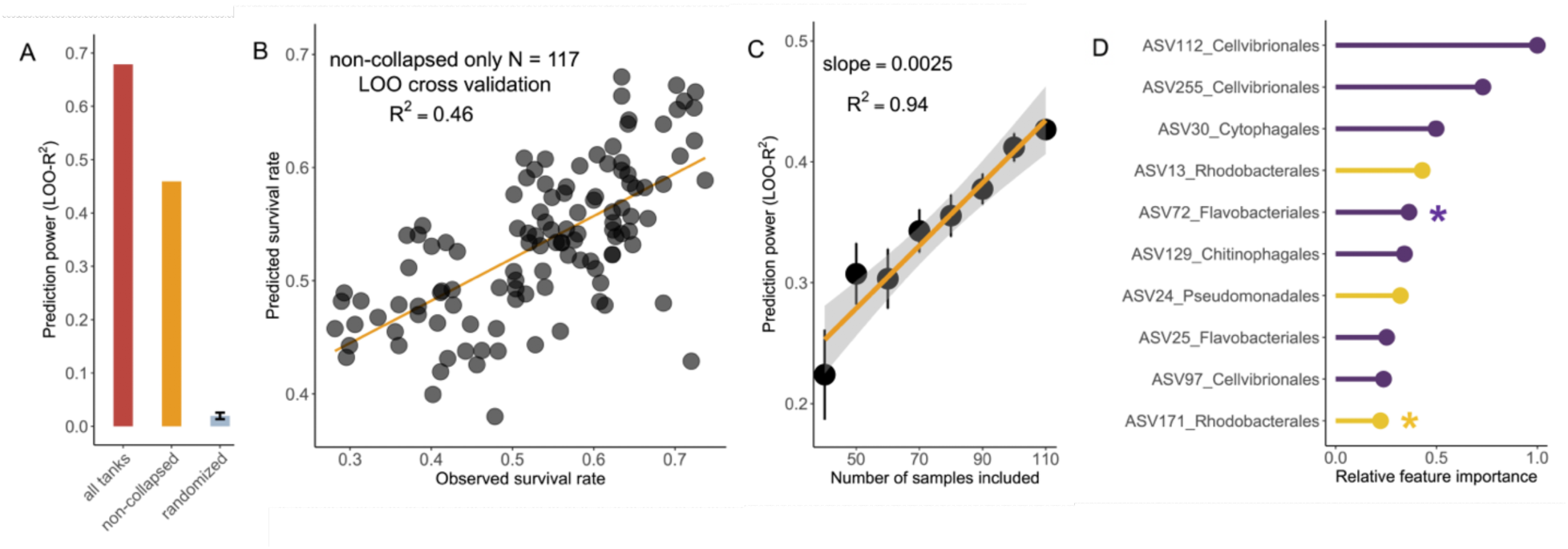
Microbiome composition as a strong predictor of shrimp larvae survival. (A) Statistical learning of microbiome composition to predict survival rate of shrimp larvae. Prediction power is indicated by leave-one-out cross-validation R^2^. (B) Observed survival rate of shrimp larvae and that predicted by microbiome composition through leave-one-out cross-validation. The 13 tanks where a bloom of *V. parahaemolyticus* led to collapse of all shrimp larvae were excluded. (C) Prediction power of microbiome composition linearly increases as the number of samples increase. On average, increasing every 10 samples leads to 2.5% increase in prediction power. (D) Taxa showing highest feature importance in statistical learning to predict survival rate of shrimp larvae. Purple lines indicate potential primary degraders. Yellow lines indicate potential scavengers.

We further seek to better understand the taxa that are important in predicting larvae survival, especially their ecological roles and genomic features. We calculated relative feature importance of each ASVs, finding that a majority of the top important ones (7 out of top 10) are from taxonomic groups well known as polysaccharide degraders^35^ (e.g., Cellvibrionales, Cytophagales and Flavobacteriales, Figure 3D). To better understand the genomic features of these important taxa, we isolated a total of 401 strains from the larvae samples, out of which 2 strains had 100% 16S rRNA gene sequence identity with the top 10 important ASVs from our Random Forest model (Figure 3D, Figure S4), one in the Rhodobacterales clade (ASV171, *Cribrihabitans* sp.) and the other in the Flavobacteriales clade (ASV72, *Psychroserpens* sp.). In addition to these two isolates, we isolated other seven strains within these two clades that are abundant (at least 1% relative abundances) in the shrimp larvae microbiome (Table S2, Figure S4), suggesting that they may have fitness advantages in the shrimp larvae-associated environment. We therefore leverage the genome of these isolates (we refer to as “abundant isolates” thereafter) to identify genes that are specifically adaptive to a shrimp-associated niche. Briefly, we filter for genes that are shared by *all* our shrimp-derived abundant isolates but are rarely present (< 10%) across non-shrimp derived close relatives (i.e., in the same genus with the shrimp-derived abundance isolates) (Figure 4A, see Methods for full details). This led to the discovery of recent horizontal gene transfer events involving these abundant shrimp larvae-derived strains in the Rhodobacterales or Flavobacteriales clades (Figure 4B, C, see Supplementary Files 1,2 for sequences). Among the Rhodobacterales strains, we observed recent horizontal transfer of genes involved in the biosynthesis of growth factors, such as vitamin B6^42^ and lysine^43^ (Figure 4B, 4C, 100% nucleotide identity between homologs in *Cribrihabitans* sp. ASV171 and *Phaeobacter italicus* ASV17), suggesting a potentially beneficial role of nutritional supplementation for the host. Among the Flavobacteriales strains, we identified a recent horizontal transfer of amidohydrolase genes, crucial for protein degradation, as well as a gene encoding a TonB-dependent receptor (TBDR) (Figure 4B, 4D, 100% nucleotide identity between homologs in *Psychroserpens* sp. ASV72 and *Meridianimaribacter flavus* ASV7). An exhaustive search of public database further revealed homologs of this TBDR, such as an isolate genome in Zhejiang Province of China (*Maribacter aurantiacus* KCTC 52409, GCF_005780245.1), as well as environmental metagenomes in Shandong Province of China (ERR2094176), both showing 100% amino acid identity with the TBDR swept across our strains isolated from the shrimp hatchery in Ecuador (Figure 4D). Both the Chinese genome and metagenome are also derived from seawater aquaculture, underscoring global ecological relevance of this gene that is specifically adaptive to aquaculture environment. Despite the functional versatility of TBDR, previous work on marine microbiology has highlighted an important role of TBDR in polysaccharide transport for Flavobacteriales^34^. Previous studies have also shown a TBDR with exactly the same domain organization (i.e., a carboxypeptidase regulatory like domain along with an outer membrane channel) functions in collagen degradation in coral-reef associated ecosystems^44^, which could inspire further biochemical studies confirming the function of this TBDR.

**Figure 4.**
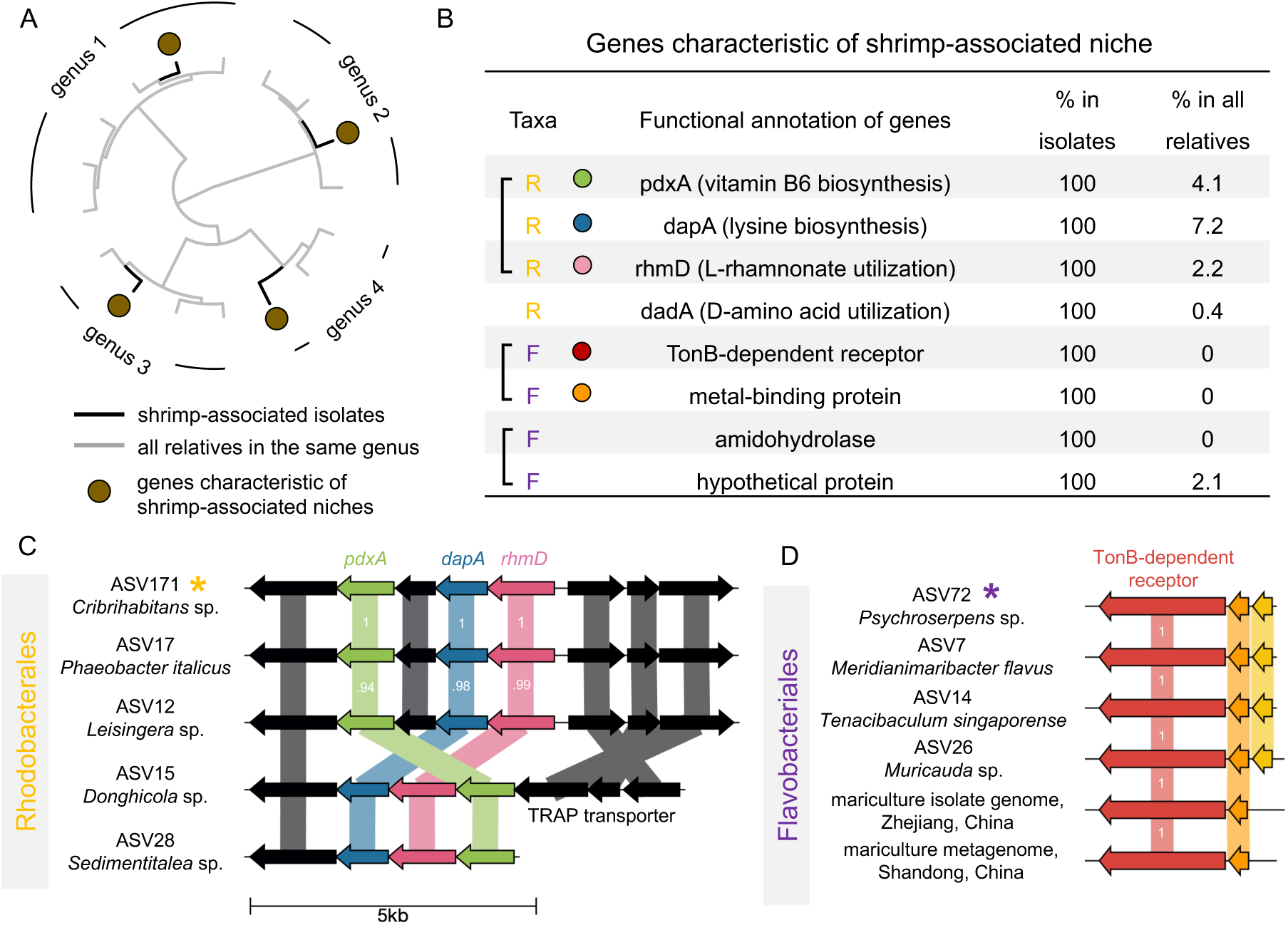
Horizontal transfer of genes that are specifically adaptive to a shrimp-associated niche. (A) Schematic illustration of comparative genomic analysis for identifying genes that are specifically adaptive of a shrimp-associated niche. The selected genes are shared by isolates that are abundant in the shrimp larvae microbiome but are present in none or very few (< 10%) of all the other publicly available genomes in the same taxonomic clades. Tips colored in black indicate strains isolated from the shrimp larvae samples. Tips colored in gray indicate publicly available genomes in the same genus with the shrimp-associated isolates. The brown circle indicates a gene that is only present in the genome of shrimp-associated isolates but not in any other genomes from the same genus. (B) A total of 8 genes are identified from the shrimp larvae-associated microbiome. Genes marked with a yellow ‘R’ are shared by all five abundant isolates in the Rhodobacterales clade. Genes marked with a purple ‘F’ are shared by all four abundant isolates in the Flavobacteriales clade. All these genes are rarely present in other genomes of the same genera from public databases (< 10%). The 16S rRNA sequences of all those nine isolates are 100% identical to ASVs in the sequenced shrimp larvae microbiome whose maximal relative abundances across tanks are at least 1%. Square brackets indicate that genes are next to each other in the genome. (C) Gene map of adjacent genes (*pdxA*, *dapA* and *rhmD*) experiencing horizontal transfer across five isolates from distinct genera in the order of Rhodobacterales. Numbers in the gene map indicates level of amino acid sequence identity between homologs. (D) Gene map of two adjacent genes (TBDR and a metal-binding protein gene) experiencing recent horizontal transfer among four isolates from distinct genera in the order of Flavobacteriales isolated from shrimp larval hatchery in Ecuador, as well as their homologs in publicly available genome and metagenome derived from aquaculture in different regions of China. All the six TBDR genes have 100% identical sequence with each other.

### Water quality is a stronger predictor of adult shrimp growth than the microbiome

Having established the significant predictive power of microbiomes for animal survival in the hatchery phase, our investigation extends to the impact of microbiome on the growth of adult shrimp in shrimp grow-out ponds. In contrast to the well-controlled conditions of hatchery tanks, adult shrimp grow in dugout ponds that extend over several hectares of water – at least 100 times larger than a hatchery tank. These are complex ecosystems that farmers manage on an individual basis to achieve maximal productivity. Despite the expected heterogeneity of these systems, we asked to what extent the microbiome of individual animals could predict the individual phenotypes, in particular, body weight. To this end, we sampled 226 adult shrimps across 6 grow-out ponds, recording the body weight of each individual. We sequenced the microbiome of the hepatopancreas for all sampled shrimps, considering that it is a primary focus of Vp infections, as well as the intestine microbiome for a subset of 76 shrimps.

The hepatopancreas data revealed a relatively simple community structure dominated by three clades: Vibrionaceae, Entomoplasmatales, and Rhizobiaceae (Figure 5A). Notably, the representative sequences for Entomoplasmatales and Rhizobiaceae differed significantly from any known organisms (92% and 87% maximum sequence identity over the 16S rRNA V3-V4 region), and likely correspond to endosymbiontic bacteria. For the Entomoplasmata, we were able to assemble a genome from metagenomic reads, which was classified as Candidatus *Hepatoplasma crinochetorum*, an endosymbiont in non-insect arthropods^45^. Again, this genome showed less than 80% average nucleotide identity with any previously sequenced organism, highlighting the presence of unknown bacterial taxa within the shrimp hepatopancreas. In contrast to the hepatopancreas microbiome, the shrimp intestine microbiome exhibits significantly greater diversity (Figure 5A, ASV richness Wilcox test *P* < 0.001). Despite the distinct compositions of the intestine and hepatopancreas microbiomes, a significant correlation exists between them (Mantel test *P* < 0.001), suggesting a systemic linkage in microbial compositions across different digestive tract sections.

**Figure 5.**
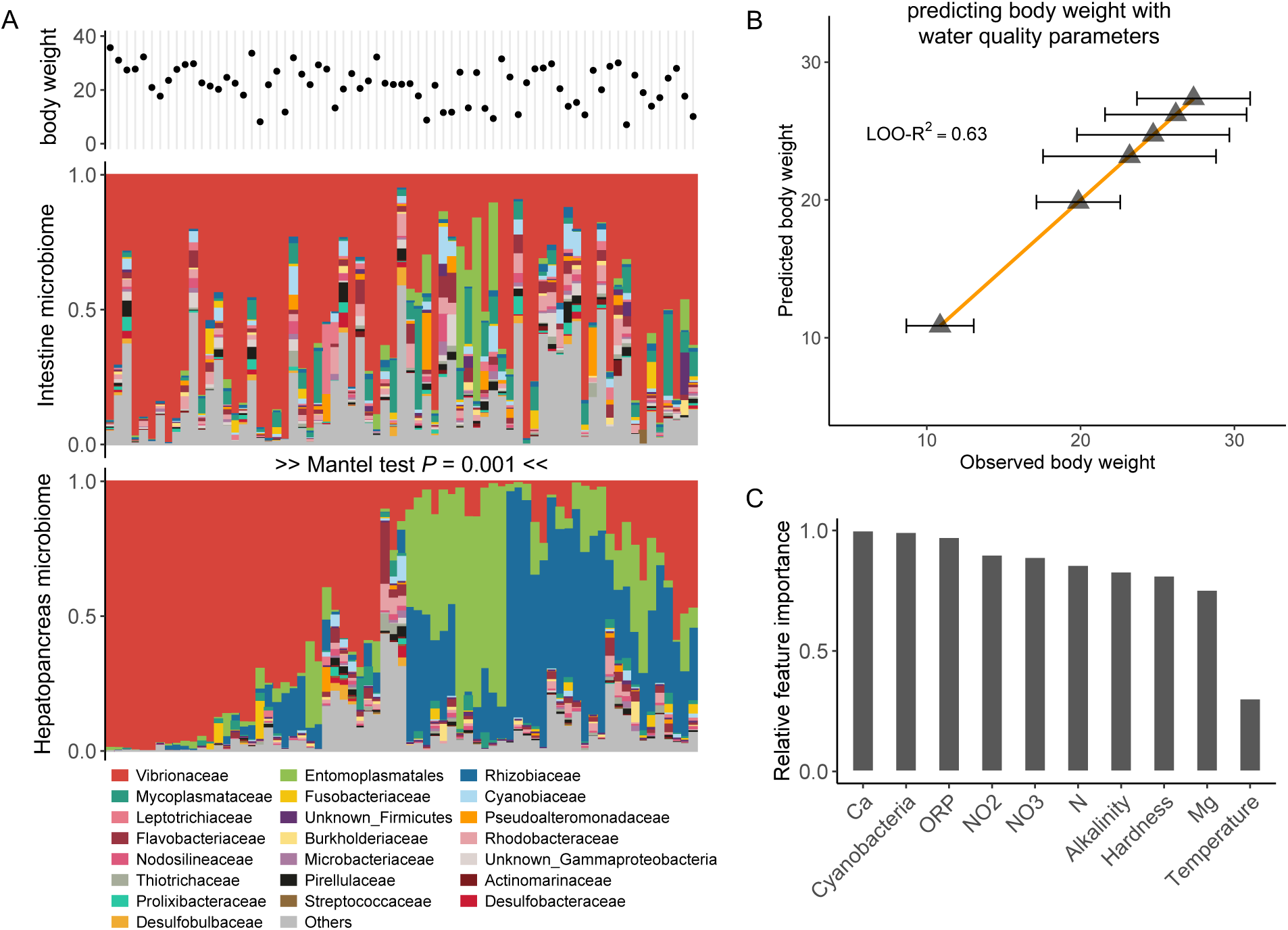
Adult shrimp body weight is better predicted by water quality than microbiome composition. (A) Adult shrimp individuals were sampled for body weight measurement and microbiome sequencing of hepatopancreas and the intestine. The microbiome composition in the intestine and that in the hepatopancreas are significantly correlated with each other. (B) Body weight of adult shrimp can be accurately predicted by water quality parameters. (C) Relative feature importance of water quality parameters in predicting adult shrimp body weight. ORP, oxidoreductive potential.

Based on this characterization of the hepatopancreas and intestine microbiomes, we sought to determine to what extent microbiome composition across the adult shrimp digestive tract could predict body weight in grow-out ponds. Our data revealed that the intestine and the hepatopancreas microbiome compositions were weak, although significant predictors of shrimp body weight (LOO-R^2^ = 0.16 for 226 samples with hepatopancreas microbiome; LOO-R^2^ = 0.18 for 76 samples with intestine microbiome). The observed R^2^ were similar to the levels observed in other livestock gut ecosystems (average cross-validated R^2^ < 0.2)^46^ and significantly different from those obtained with randomized data (LOO-R^2^ = 0.03 ± 0.01 for 100 random tests, *P* < 0.001). In contrast to the high degree of unexplained variance across individuals, our analysis of the metadata showed that the average body weight of individual ponds correlates strongly with water quality parameters (LOO-R^2^ = 0.63 for 226 samples, Figure 5B), indicating that the average animal weight is primarily controlled by water chemistry. Among the key factors revealed by the data alkalinity, hardness, oxidoreductive potential, nitrogen concentration, and cyanobacteria density emerge as key predictors (Figure 5C). Alkalinity, indicating buffer capacity of the pond water, has been shown to closely correlated productivity in shrimp farms^47^. Hardness is determined by the concentration of dissolved calcium and magnesium, which are important for exoskeleton development and osmoregulation^47,48^. Cyanobacterial blooms can have adverse effects due to toxin production and eutrophication^49^. These factors highlight the complex dynamics between water chemistry, eutrophication and animal health that control productivity, and thus feed-efficiency, in large-scale aquaculture facilities.

## Discussion

In this study, we discovered that the microbiome composition alone accounts for 50% of shrimp larvae survival, highlighting the potential of microbiome as an early indicator for host health.

The microbiome associated with these larvae likely affects host production by aiding in the breakdown of polymeric organic materials or by synthesizing beneficial growth factors such as vitamins, as suggested by our comparative genomics analysis. While prior research has demonstrated that mapping the structure to the function of microbial communities can facilitate functional predictions^50,51^, these studies were primarily based on simplified microcosms in laboratory settings. Our investigation extends these findings by establishing a robust structure-function relationship within a complex, real-world ecosystem. Notably, our study reveals that tree-based models, like Random Forest and Gradient Boosting, largely outperform linear models in predicting functional output in this intricate real-world context (Figure S5). This suggests that the functional landscape of natural microbiomes might be more complex than that synthetic microbial communities with low biodiversity.

With respect to adult shrimps, which develop in less-controlled, complex ecosystems like grow-out ponds, our analysis shows that water quality parameters are strong predictors of the average shrimp weight per pond. However, there is a large degree of variance in individual body weight, which the microbiome can only partially explain (< 20%) likely due to variable chemical and biological conditions in these environments. This suggests that hatcheries may be a better target to design microbiome-based interventions during early developmental stages, especially considering that, as shown in our data, currently used probiotics based on non-native bacteria display low engraftment rates.

Our study uncovers an understudied taxonomic and functional diversity within the shrimp-associated microbiome, opening avenues for microbiome engineering. For example, a gene cluster involved in cofactor biosynthesis, prevalent across five genera of Rhodobacterales, shows strong signs of adaptive potential. Similarly, the widespread horizontal transfer of the TBDR gene among our isolates in Ecuador, as well as those from Asian aquacultures, suggests a universal fitness advantage for various strains of Flavobacteriales in widely distributed aquaculture systems. This points to the potential of engineering polysaccharide degradation-assisting probiotics (e.g., Flavobacteriales) that are adaptable across global shrimp aquaculture settings. These findings provide promising targets for downstream genetic and biochemical studies, paving the way for innovative strategies to optimize shrimp production by microbiome engineering.

## Methods

### Sampling in the hatchery

Microbiome sampling was performed within commercial hatchery at the coast of Ecuador from January 27 to February 19, 2021. A total of 130 tanks with *P. vannamei* was sampled at the end of the production cycle, when the survival rate of shrimp larvae was also recorded. There were 13 tanks presenting an outbreak disease and samples of larvae of these tanks were collected.

Furthermore, for a subset of 4 tanks, we sampled the shrimp larvae every two days throughout the developmental stages from nauplius (N), zoea (Z), mysis (M) and postlarvae (PL). This results in 9 timepoints spanning 18 days from N5 to PL10 stages for each tank. The shrimp larvae were fed with microalgae by the zoea stage (N5 and Z2 data points in our samples), and then *Artemia* since the mysis stage (M1 to PL10 data points in our samples). The feed microalgae, *Artemia*, together with the ambient seawater and administrated probiotics were also sampled for microbiome sequencing to track the potential source of microbiome in the shrimp larvae.

### Sampling in shrimp grow-out ponds

A total of 224 adult shrimp were collected from six different pools from a large-scale shrimp farm located in the coast of Ecuador and stored in cryovials with DMSO. The work was carried out wearing KN95 masks and nitrile gloves in a sterile area to prevent contamination. Adult shrimps were collected before feeding began, starting at 7 AM and finishing by 10 AM. Animals were kept in buckets with aeration and water from their respective pools until their turn for collection. Each animal was weighed and photographed, after which they were taken to a sterile area for dissection of the intestine and the hepatopancreas. Half of the hepatopancreas was used for DNA sequencing and the other half for microscopic observation, to record the degree of lipid vacuole fullness and changes in color that could indicate sickness. The remaining half of the hepatopancreas and intestine were immediately frozen in separate cryovials with liquid nitrogen and kept at -20°C until shipment.

Water quality was analyzed by measuring pH, salinity, temperature, total ammonium nitrogen, nitrite, nitrate, total nitrogen, phosphate, phosphorus, N:P ratio, alkalinity, calcium, calcium hardness, magnesium, magnesium hardness, potassium, Ca:Mg:K ratio, hydrogen sulfide (H2S), Redox potential and turbidity. Algae type quantification and identification were also performed by counting in a Neubauer chamber.

### DNA extraction and amplicon sequencing

DNA extraction of the shrimp larvae was performed with Gentra Puregene Tissue Kit (QIAGEN) following the protocol provided by the manufacturer, except that bead beating was used to increase the extraction efficiency for 60 seconds at 5000 rpm. In brief, the extraction procedure includes cell lysis (1:1 cell lysis solution), RNA removal (4μL of RNAse A), protein precipitation (250 μL Protein Precipitation Solution), DNA precipitation (100% isopropanol) and purification (70% ethanol). Purified DNA was shipped to Argonne National Laboratory (Lemont, IL) for amplicon sequencing on an Illumina MiSeq targeting the V4 region of 16S rRNA using the 515F and 806R primers. Sequence-specific Peptide nucleic acid (PNA) clamps were used to block the amplification of host-derived mitochondrial 16S sequences at V4 region.

Shrimp hepatopancreas and intestine samples were torn into small pieces using tweezers. These fragments were further homogenized using a TissueLyser III (Qiagen) for 5 minutes at 25 Hz with PowerBeads Pro plates (Qiagen). Following this, DNA extraction was carried out on the resulting turbid liquid using the DNAdvance Kit (Beckman Coulter), according to the manufacturer’s protocol. Briefly, the extraction process included cell lysis using a 1:1 cell lysis solution, DNA binding to magnetic beads, and purification with 70% ethanol. Amplicon sequencing were performed at Seqcenter (Pittsburg, PA) with Zymo Quick-16S Plus Library Prep methods to amplify the V3/V4 region with 341F and 806R primers. Illumina sequencing was performed on an Illumina NovaSeq X Plus sequencer in one or more multiplexed shared-flow-cell runs, producing 2x151bp paired-end reads.

Amplicon sequences analyses were performed on the MIT Engaging computing cluster, where we used QIIME2 v2019.2^52^ to demultiplex the raw reads, and DADA2 plugin^53^ to generate Amplicon Sequence Variants (ASVs). Taxonomy of representative ASVs were assigned with the classify-sklearn method by QIIME2 v2019.2.

### Strain isolation and genotyping

We first isolated ∼200 isolates using Difco^TM^ Marine Broth 2216 media (BD). Colonies were isolated by first growing on agar plates prepared with MBL medium^54^ containing homogenized Artemia as the sole carbon source, and then streaking to single colony purity on agar plates prepared with 1/10 diluted Marine Broth 2216 medium (adjusted for salt) containing 0.2% N-acetyl-glucosamine as the sole carbon source. A total of ∼ 200 Rhodobacterales strains were further isolated with a customized media recipe favoring growth of Rhodobacterales species among other marine copiotrophic heterotrophs^35^. The media is MBL minimal media with 40mM sodium succinate added as the carbon source. Marine Broth 2216 was spiked into the media with a dilution factor of 1:80 to facilitate bacterial growth. Vp was selectively isolated by the CHROMagar^TM^ *Vibrio* (CaV) selective media, where the mauve colonies are picked for genotyping. Genotyping of all isolates were performed by full-length 16S Sanger sequencing (GENEWIZ at Azenta Life Sciences). 16S sequences of isolates were aligned to ASV sequences from the shrimp larvae mirobiome with blastn in the blast suite v2.12.0. A maximal-likelihood phylogenetic tree of representative 16S sequences was constructed with MEGA v11.

### Whole genome sequencing and genomic data processing

Shotgun whole genome sequencing for representative isolates of Flavobacteriales and Rhodobacterales were performed at the SeqCenter (Pittsburgh, PA), on an Illumina NextSeq 2000 platform (2x151bp pair-ended). Raw reads were trimmed to remove adaptors and low-quality bases (-m pe -q 20) with Skewer v0.2.2^55^. The remaining paired reads were checked for quality with FastQC v0.11.9. Quality-filtered reads were assembled into contigs with MEGAHIT v1.2.9^56^.

Nanopore long-read sequencing was combined with shotgun whole genome sequencing also at the SeqCenter (Pittsburgh, PA) to close the genome of Vp. Closed genome was assembled using Unicycler v0.4.9^57^ by combining Illumina short reads and Nanopore long reads, resulting in 2 chromosomes and 3 circular plasmids. The assembly graph in gfa format was visualized by Bandage v0.8.1^58^. Coding sequences are predicted by prodigal v2.6.3, followed by functional annotation by eggnog-mapper v2^59^ (--go_evidence non-electronic --target_orthologs all --seed_ortholog_evalue 0.001 --seed_ortholog_score 60).

### Metagenomic sequencing and processing

We did shotgun metagenomic sequencing for 6 shrimp hepatopancreas samples (2 samples with highest relative abundance of Vibrionaceae, 2 samples with highest relative abundance of Entomoplasmatales and 2 samples with highest relative abundance of Rhizobiaceae). Shotgun sequencing including library preparation was performed at SeqCenter (Pittsburgh, PA) on an Illumina platform. We performed quality check for pair-ended fastq files with FastQC v0.11.9, after which low-quality bases were trimmed by Skewer v0.2.2 ^55^ (-m pe -q 20). Quality-filtered reads were first filtered with Bowtie2 v2.5.3 (--un-conc) to get rid of all the sequence aligned with the *P. vannamei* genome (NCBI RefSeq accession GCF_003789085). All the unaligned sequence was then used to assemble contigs with MEGAHIT v1.2.9. The reads that were unaligned to the *P. vannamei* genome were mapped back to assembled contigs with Bowtie2^60^2.5.3 to generate coverage table. Metagenomic assembled genomes (MAGs) were generated with CONCOCT^61^ v1.1.0 (-c 10000). Generated bins were evaluated for completeness and contamination using CheckM^62^ v1.1.2. Taxonomic classification of MAGs was performed by GTDB-tk v2.1.0^63^. We were able to generate a MAG of *Vibrio fischeri* (completeness 100%, contamination 0.5%) and a MAG of *Hepatoplasma crinochetorum* (completeness 87.5%, contamination 0%) through this effort.

### Identifying core clades from global shrimp-associated microbiome

We compiled a comprehensive global dataset of amplicon microbiome sequences, specifically targeting the 16S rRNA V4 region with an approximate overlap of ∼250 base pairs (bp). This overlap facilitates the downstream analysis of core microbial clades. We retrieved amplicon sequencing data in FASTQ format from the National Center for Biotechnology Information

(NCBI) database. Using DADA2^53^ within QIIME2^52^ (version 2019.2), we processed samples from various studies, yielding 48,514 shrimp-associated amplicon sequence variants (ASVs) and 7,999 coastal seawater ASVs.

For sequence alignment, we utilized MAFFT^64^ (version 7.245), followed by sequence trimming to uniform lengths using TrimAl^65^. To ensure high-quality alignment of the vast number of sequences, we initially conducted stringent alignments (--maxiterate 1000) focusing on ASVs from the most abundant taxonomic groups, including Vibrionales, Rhodobacterales, Flavobacteriales, and Alteromonadales. These aligned sequences were trimmed to uniform lengths, creating a ’backbone’ alignment. Subsequent sequences were integrated into this backbone using MAFFT’s --add function. We then merged this composite alignment with pre-aligned reference sequences from the SILVA database^66^ (release 132, Nr99) to construct phylogenetic trees for each major clade. After a final trim with Trimal, the sequences were standardized to approximately 250 bp. We constructed the phylogenetic trees using FastTreeMP^67^, employing the -nt, -gtr, and -gamma options, and visualized the trees with iTOL^68^. Moreover, we conducted homologous clustering of all shrimp-derived sequences using MMseqs2 (-s 7.5 –min-seq-id 1)^69^. This analysis supports the power-law distribution illustrated in Figure 1D.

To identify core shrimp-associated clades, we applied three stringent criteria. First, a clade must show significant enrichment (adjusted P < 0.05, with Bonferroni correction) of shrimp-derived sequences, assuming a binomial distribution of these sequences across the entire phylogenetic tree. Second, the clade must encompass shrimp-associated sequences that are globally ubiquitous, present in at least 80% of sampled global shrimp cultures. Third, we required complete homology within each clade, selecting only those with zero branch length, indicating 100% sequence identity. Ultimately, five clades met these criteria for significant enrichment, global prevalence, and stringent homology. We assigned representative taxonomy to each clade based on the dominant taxonomy of the corresponding SILVA^66^ reference sequences. For the overarching phylogenetic analysis, we constructed a tree with all 56,513 ASV sequences derived from both shrimp and coastal seawater. We labeled each tree tip according to its source. The observed mean phylogenetic distance (MPD) between shrimp-associated and coastal seawater sequences was calculated. To assess the significance of our findings, we randomized the MPD calculations 1,000 times by reshuffling the tip labels. The Net Relatedness Index (NRI) is calculated by the standardized effect size of the MPD, which is (𝑜𝑀𝑃𝐷 − 𝑚𝑒𝑎𝑛(𝑟𝑀𝑃𝐷))⁄𝑠𝑑(𝑟𝑀𝑃𝐷).

### Identifying characteristic genes of shrimp-associated niche

We aligned the 16S rRNA gene sequences of our 501 isolates to the Flavobacteriales and Rhodobacterales ASV sequences in the microbiome. This resulted in 4 Flavobacteriales isolates and 5 Rhodobacterales isolates with 100% identical sequences with an *abundant* ASV in the shrimp-larvae microbiome (at least 1% relative abundance in any sample). These nine isolates spanned nine distinct genera, prompting us to retrieve all available genomes for these genera from the RefSeq database. This search yielded 394 genomes for Flavobacteriales and 222 for Rhodobacterales.

Subsequently, we conducted protein sequence clustering using MMseqs2^69^ (-s 7.5 -min-seq-id 0.5 -c 0.5). This analysis produced a matrix, organizing proteins by rows and genomes by columns. We then refined our search to identify proteins universally present in our shrimp larvae isolates yet scarcely found (< 10%) within other genomes of the same genera. This rigorous filtering process highlighted eight genes meeting these specific criteria. We utilized EggNOG-mapper^59^ v2 for functional annotation the identified protein sequences. To visualize gene maps, we employed Clincker^70^, facilitating a comprehensive representation of the gene distribution.

Moreover, our extensive search within the public metagenomic database MGnify^71^ led to the discovery of a 100% homologous sequence in mariculture environments in China.

### Machine learning-based predictions

We implemented a Random Forest (RF) model to predict outcomes like shrimp larvae survival rate and adult shrimp body weight based on microbiome composition, using leave-one-out (LOO) cross-validation for evaluation. In this method, the RF model, developed with the R package randomForest^72^, is trained on all but one sample to predict the outcome for the excluded sample, a process repeated for each sample. Predictive accuracy is quantified by comparing observed and predicted values through R² from linear regression, with robustness checked against 100 randomized trials of the outcome vector. To assess how sample size affects prediction power, we varied sample subsets (n= 40 to 110, in increments of 10) and conducted LOO cross-validation 20 times for each size. We also compared RF performance against other machine learning methods like gradient boosting and lasso regression, to gauge their efficacy (as detailed in Figure S5), providing a comprehensive view of the predictive capabilities of various approaches within our dataset.

### Other statistical analysis

All other statistical analyses are implemented with R^73^ v4.1.3. World map is visualized using the default world map in R package ggmap^74^ v3.0.0.

## Data availability

All raw data generated by this study has been deposited on NCBI under BioProject PRJNA1144440. This BioProject includes (1) amplicon sequencing data of shrimp larvae including its associated feed and environmental samples, (2) amplicon sequencing data of hepatopancreas and intestine samples of adult shrimp, (3) shotgun metagenomic sequencing data of selected adult shrimp hepatopancreas samples, (4) draft genomes assembled for the 9 abundant isolates shown in Figure 4. The compiled global aquaculture shrimp microbiome dataset is publicly available at 10.5281/zenodo.13241422 for free use. Detailed information and sequences of the horizontal transferred genes are provided as supplementary file 1 and 2.

## Acknowledgement

We thank Salvador Almagro-Moreno, and members of the Cordero lab, in particular Andreas Sichert, Rachel Gregor, Gabriel Vercelli and Suryateja Jammalamadaka, for helpful discussions and valuable assistance. X.S thanks Zhijian Huang, Shenzheng Zeng, Haipeng Guo and Mengmeng Wang for their help in collecting the global microbiome datasets. X.S also thanks Shiwei Wang and Victoria Marando in Laura Kiessling lab at MIT for their help on understanding the interactions between aquaculture isolates.

## Notes

### Competing Interest Statement

The authors have declared no competing interest.

### Summary of Updates

Removed an excessive symbol in the title

