## supplementary materials for "Microbiome determinants of productivity in whiteleg shrimp aquaculture"

**File S1** Genomic region containing pdxA, dapA and rhmD in ASV171 Cribrihabitans sp. (Separate file attached).

**File S2** Genomic region containing TonB-dependent receptor in ASV72 Psychroserpens sp. (Separate file attached).

**Table S1** List of microbiome studies contributing to our compilation of the global shrimp-associated microbiome.

| Site | #Samples | System | Type | Reference |
|---|---|---|---|---|
| Dongying (Cdy) | 2 | shrimp | instestine | 18 |
| Ningbo (Cnb) | 121 | shrimp | instestine | 19 |
| Wenzhou (Cwz) | 6 | shrimp | instestine | 20 |
| Zhuhai (Czh) | 75 | shrimp | instestine | 21 |
| Maoming (Cmm) | 41 | shrimp | instestine | 22 |
| Wenchang (Cwc) | 101 | shrimp | larvae | 23 |
| Vietnam (Vt) | 10 | shrimp | instestine | 24 |
| Thailand (Th) | 48 | shrimp | instestine | 25 |
| Malaysia (Ma) | 48 | shrimp | instestine | 24 |
| Mexico (Me) | 23 | shrimp | instestine | 26 |
| Ecuador (Ec) | 131 | shrimp | larvae | this study |
| Brazil (Br) | 4 | shrimp | instestine | 27 |
| SW1 | 95 | seawater | seawater | 28 |
| SW2 | 47 | seawater | seawater | 30 |
| SW3 | 12 | seawater | seawater | 29 |

**Table S2** The nine isolates from the clade of Flavobacteriales and the clade of Rhodobacterales whose corresponding ASV (100% sequence identity) are abundant in the original shrimp larvae-associated microbiome.

| | GDTB taxonomy | maximal relative abundance |
|---|---|---|
| ASV7_Flavobacteriales | *Meridianimaribacter flavus* | 0.24 |
| ASV14_Flavobacteriales | *Tenacibaculum singaporense* | 0.09 |
| ASV26_Flavobacteriales | *Muricauda* sp. | 0.03 |
| ASV72_Flavobacteriales | *Psychroserpens* sp. | 0.16 |
| ASV12_Rhodobacterales | *Leisingera* sp. | 0.24 |
| ASV15_Rhodobacterales | *Donghicola* sp. | 0.14 |
| ASV17_Rhodobacterales | *Phaeobacter italicus* | 0.05 |
| ASV28_Rhodobacterales | *Sedimentitalea* sp. | 0.07 |
| ASV171_Rhodobacterales | *Cribrihabitans* sp. | 0.01 |

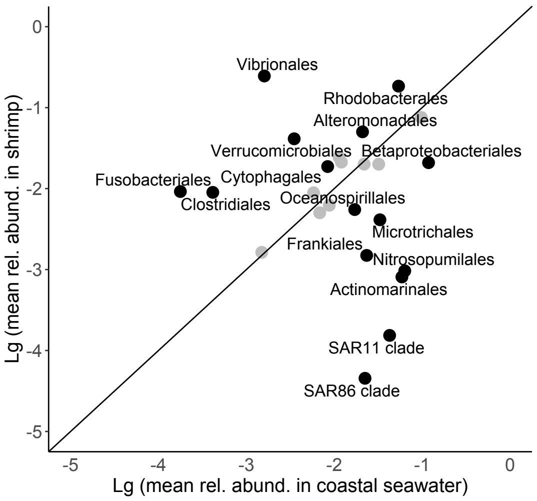


**Figure S1** Enrichment of microbial taxa in coastal seawater microbiome and shrimp-associated microbiome. Black dots indicate significant enrichment in either shrimp-associated or coastal seawater microbiome (fold change > 2 and adjusted P value < 0.05).

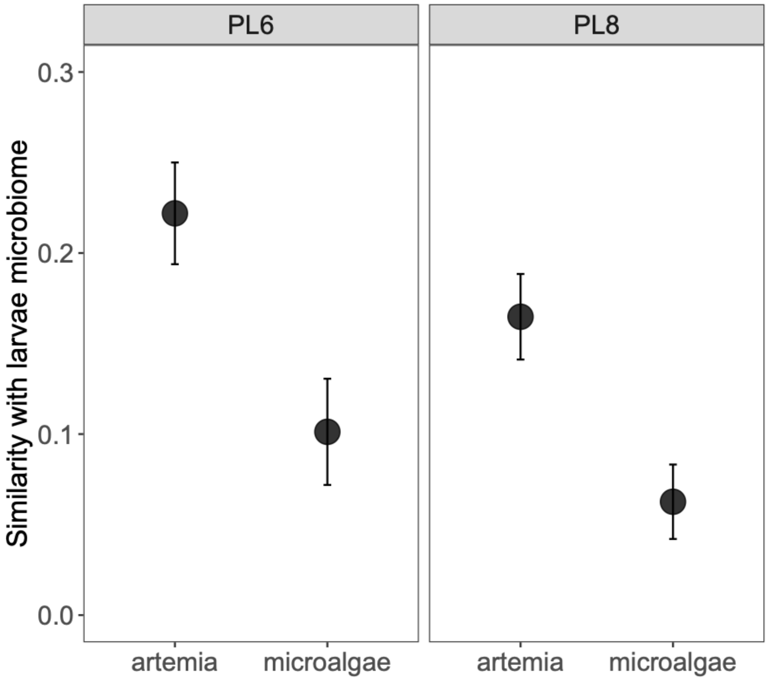


**Figure S2** At PL6 and PL8 stages, feed artemia has larger impact on the larvae microbiome composition than feed microalgae. All artemia and microalgae samples that are fed before PL6 or PL8 stages were used to calculate similarity with the microbiome composition of shrimp larvae microbiome at the respective stage.

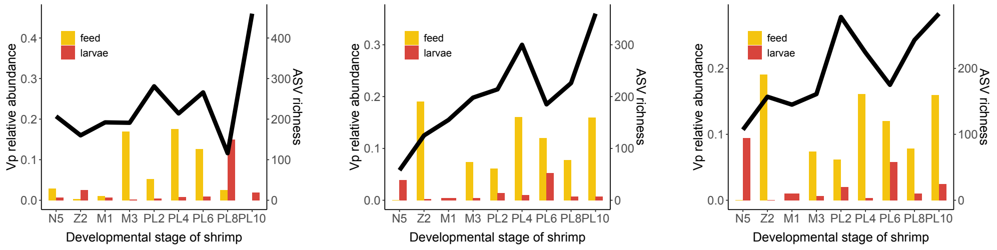


**Figure S3** Relative abundance of Vp in larvae and feed microbiome in other three tanks where we temporally sampled across the developmental stages. Black line indicates richness of shrimp larvae-associated microbiome across developmental stages. Red bar and yellow bar indicate relative abundances of Vp across developmental stages in shrimp larvae microbiome and microbiome in the feed.

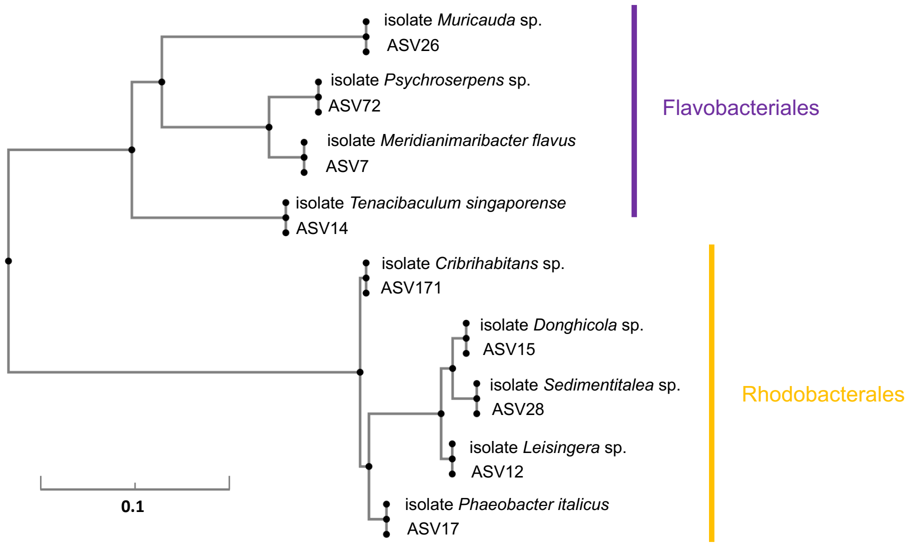


**Figure S4** Phylogenetic tree of 9 isolates in Flavobacteriales and Rhodobacterales with 100% identical sequences with an *abundant* ASV in the shrimp-larvae microbiome (at least 1% relative abundance in any tanks).

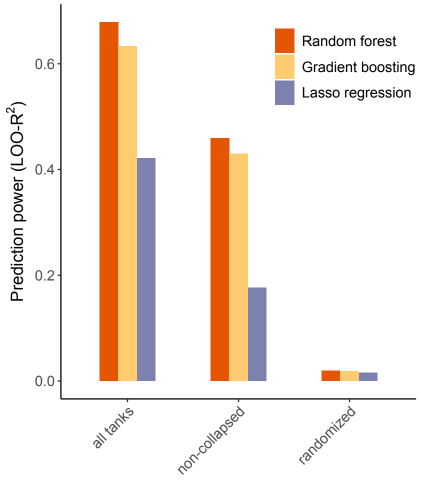


**Figure S5** Predicting shrimp larvae survival rate by microbiome composition, using different models (Random Forest, Gradient Boosting and LASSO regression).
